# Acute effect of antiseizure drugs on background oscillations in *Scn1a*^A1783V^ Dravet syndrome mouse model

**DOI:** 10.1101/2022.11.29.518351

**Authors:** Shir Quinn, Marina Brusel, Mor Ovadia, Moran Rubinstein

## Abstract

**Objective:** Dravet syndrome (Dravet) is a rare and severe form of developmental epileptic encephalopathy. First-line treatment for DS patients includes valproic acid (VA) or clobazam with or without stiripentol (CLB+STP), while sodium channel blockers like carbamazepine (CBZ) or lamotrigine (LTG) are contraindicated. As patients are rarely seizure-free, drug therapy focuses on reducing the seizure burden, as reported by caregivers. In addition to their effect on epileptic phenotypes, antiseizure medications (ASMs) were shown to modify the properties of background neuronal activity. Nevertheless, little is known about these background properties alternations in Dravet.

**Methods:** Utilizing Dravet mice (DS, *Scn1a*^A1783V/WT^), we tested the acute effect of several ASMs on background electrocorticography (ECoG) activity and frequency of interictal spikes.

**Results:** Compared to wild-type mice, background ECoG activity in DS had lower power and reduced phase coherence, which was not corrected by any of the tested ASMs. However, acute administration of Dravet-recommended drugs, including VA or a combination of CLB+STP, caused, in most mice, a reduction of frequency of interictal spikes, alongside an increase in the relative contribution of the beta frequency band. Conversely, CBZ and LTG increased the frequency of interictal spikes with no effect on background spectral properties. Moreover, we uncovered a correlation between the reduction in interictal spike frequency, the drug-induced effect on the power of background activity, and a spectral shift toward higher frequency bands.

**Significance:** These data provide a comprehensive analysis of the effect of selected ASMs on the properties of background neuronal oscillations and highlight a possible correlation between their effect on epilepsy and background activity. Thus, examining these properties, following an acute administration, may be used as an additional tool for rapid evaluation of the therapeutic potential of ASMs.

**Key Points:**

- Reduced background power and phase coherence in Dravet mice
- DS-recommended medicines (VA, CLB+STP) increase the relative beta power
- DS-contraindicated drugs (CBZ, LTG) do not cause spectral changes
- Correlation between reduction in background power and interictal spike frequency
- Correlation between theta to beta bands ratio and interictal spike frequency

## Introduction

Dravet syndrome (Dravet) is a rare and severe form of developmental epileptic encephalopathy (DEE). Most cases are caused by heterozygous de novo mutations in the *SCN1A* gene, encoding the alpha subunit of the voltage-gated sodium channel type I (Na_V_1.1). The first sign of the disease is febrile seizures that soon progress to refractory spontaneous seizures with developmental delays^1,2^. Electroencephalogram (EEG) background activity is typically normal at the onset of seizures, but interictal epileptic discharges can be observed ^3–5^.

Dravet seizures are difficult to control, even with polytherapy. Drug treatment includes valproic acid (VA) or clobazam (CLB) as first-line drugs, with stiripentol (STP), fenfluramine, or cannabidiol as a second-line add-on treatment. Conversely, sodium channel blockers such as carbamazepine (CBZ) or lamotrigine (LTG) are contraindicated, as these can aggravate the seizures^5^.

Dravet mouse (DS mice) models are an exceptionally good genocopy and phenocopy of the human syndrome. DS mice are mostly asymptomatic until their fourth week of life (postnatal day (P) 20-27), when they begin exhibiting spontaneous seizures and profound premature mortality ^6–10^. Electrocorticography (ECoG) recordings from DS mice at their fourth week of life demonstrated an unaltered spectral profile, similar to Dravet patients ^11–13^.

However, analyses of the non-normalized power spectral density (PSD) in naïve, unmedicated mice showed reduced power. Moreover, mice that died prematurely had the lowest power ^12,13^. Considering this link between background oscillations and epilepsy in DS mice, we set out to test the effect of several antiseizure medications (ASMs) on ECoG spectral properties. The effect of ASMs on background EEG activity was studied before, but not in Dravet. VA was shown to reduce the power of background EEG in patients with idiopathic generalized epilepsy, and juvenile myoclonic epilepsy ^14,15^, and CLB caused spectral changes in rabbits ^16^ and kindled rats ^17^.

Utilizing DS mice (*Scn1a*^A1783V/WT^) we show that acute administration of VA, a combination of CLB+STP, or STP alone, was associated with a reduction in the frequency of interictal spikes in most of the tested mice, along with a significant increase in relative beta band contribution to the total power.

Conversely, CBZ and LTG, contraindicated in Dravet, increased the frequency of interictal spikes with no effect on the spectral properties of background activity. Moreover, we uncovered a correlation between the reduction in interictal spike frequency and the redistribution of power towards increased beta and gamma band contribution and reduced theta band. Together, these data demonstrate that ASMs modulate background neuronal activity and that these modulations may correlate with their effect on seizure control. Thus, we propose that performing such measurements in conjunction with acute drug administration may be used to predict the therapeutic potential.

## Methods

### Animals and Surgery

All animal experiments were approved by the Institutional Animal Care and Use Committee (IACUC) of Tel Aviv University. Mice used in this study were housed in a standard animal facility at the Goldschleger Eye Institute at a constant (22°C) temperature, on 12-hour light/dark cycles, with ad libitum access to food and water.

DS mice harboring the global *Scn1a*^A1783V/WT^ mutation on pure C57BL/6J background were generated as described before ^12,18^. Electrode implantation was done on P19-P25, as previously described ^12^. Mice were allowed to recover for at least 48 hours before recording. (See Supplementary material for more details).

### Drugs administration

The drugs that were used were: valproic acid (VA, 300 mg/kg in saline; Sigma-Aldrich), stiripentol (STP, 150mg/kg in sesame oil; EDQM), a combination of clobazam and stiripentol (CLB+STP, 5 mg/kg + 100 mg/kg in sesame oil, respectively; Angene Chemical, EDQM), carbamazepine (CBZ, 20 mg/kg in 30% polyethylene glycol 400; Alomone Labs) and lamotrigine (LTG, 10 mg/kg in 30% polyethylene glycol 400; Alomone Labs). Drug dosages were chosen to reach therapeutic-relevant concentrations following acute administration ^19^. All drugs were administered as a single IP injection in a volume of 10 ml/kg.

### Data acquisition and ECoG signal pre-processing

Video-ECoG recordings were obtained from freely behaving mice on their P21-P27 ^12^. After two hours of recording, one of the drugs was administered. The EMG signal was denoised and smoothed using a custom-written Python script based on methods described before ^20^. Power spectral densities were computed for each epoch using Welch’s method with a 50% overlap Hann window. A custom-written threshold algorithm was used to extract artifact-free 5-second epochs, in which the mice were awake and not moving. All automatically selected epochs were concatenated and manually inspected. For each mouse, at least 100 epochs were analyzed. To quantify phase synchrony between the left and right somatosensory electrodes, we calculated the interhemispheric coherence coefficient (coherence) using a custom-written Python script ^21^. The frequency of interictal spikes was quantified using the spike histogram module in LabChart 8 (ADInstruments), with a threshold of 4-5 times the standard deviation and 250 ms maximal duration. The spikes were then manually inspected. (See Supplementary material for more details).

### Thermally-induced seizures

Thermal induction of seizures was performed as previously described ^18^. Drugs that were used for assessing thermal induction (TI) of seizures were: *i*) VA (300 mg/kg); *ii*) CLB+STP (5 mg/kg + 100 mg/kg). Control DS mice were injected with the appropriate vehicle control.

### Statistics

Statistical analyses were performed using Prism 9 (GraphPad Software). Data are reported as mean ± SE. Statistical analysis utilized Student’s t-test or the non-parametric Mann-Whitney test (for data with normal distribution or data that did not distribute normally). To compare the effectiveness of drugs we used the parametric paired Student’s t-test or the Wilcoxon matched-pairs signed rank test (for data with normal distribution or data that did not distribute normally). To compare the effect on normalized power spectral density (PSD) or coherence, we used Two-Way repeated measures ANOVA followed by Sidak posthoc analysis. For the correlation analysis, the Spearman rank-order correlation coefficient was used. Statistical analysis of pie charts was performed using Fisher’s exact test. We considered p < 0.05 as statistically significant.

## Results

### Reduced background ECoG power and interhemispheric coherence in DS mice

Global neuronal activity in juvenile WT and DS littermates (P21-P27) was examined using simultaneous video-ECoG recordings, followed by quantitative analyses of background activity spectral properties In accord with previous reports ^12,13^, a global reduction in total power was seen in DS mice at this age, observed in all frequency bands between 1 to 30 Hz (Figure 1A-D). Extracting the relative power for each band revealed a higher contribution of the gamma band (30 - 100 Hz) and a reduced contribution of theta (4 - 8 Hz) and alpha bands (8 - 12 Hz) (Figure 1E).

**Figure. 1.**
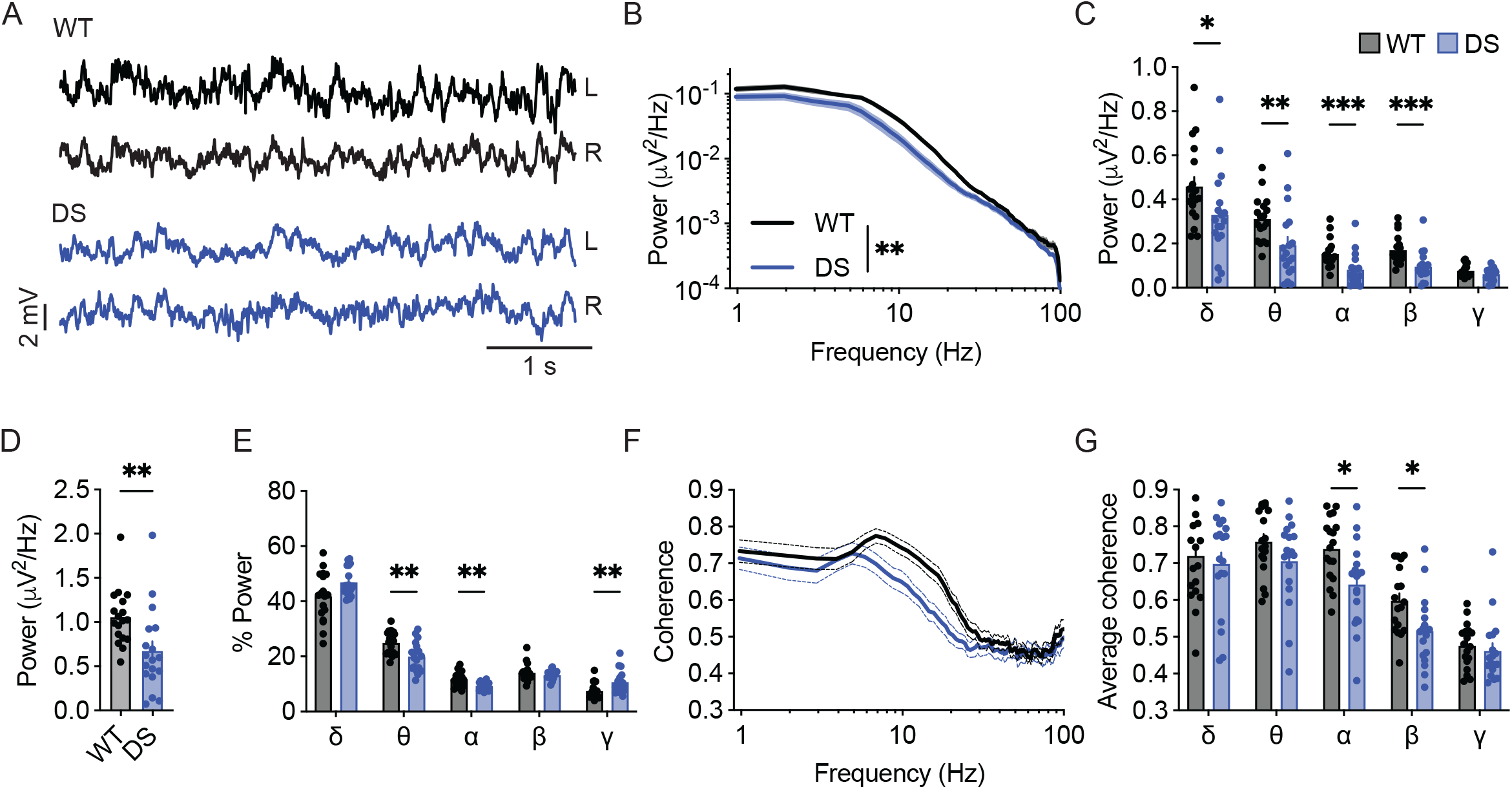
Reduced power of background activity and interhemispheric coherence in DS mice. **(A)** Representative traces of background ECoGs in WT and DS mice. **(B)** ECoG power density profiles. **(C)** Total power for the following frequency bands: δ, 1-3.9 Hz; θ, 4.8-7.8 Hz; α, 8.7-11.7Hz; β, 12.6-29.3 Hz; γ, 30.2-100 Hz. **(D**) Total power. **(E)** The relative power (%) of each frequency band. **(F)** Interhemispheric coherence plotted over the 1-100 Hz spectrum. **(G)** The mean interhemispheric coherence for each frequency band. WT, n = 18; DS, n = 18. *p < 0.05, **p< 0.01, ***p < 0.001.

Altered interhemispheric phase synchrony, or coherence, was found in various brain disorders, including epilepsy and autism ^22^. Here, we found reduced coherence in DS mice compared to WT controls, mainly in the alpha and beta frequency bands (Figure 1F-G), indicating deficits in synchrony between frequency components of background ECoG. Together, quantitative analyses of background ECoG recordings from DS mice demonstrated reduced power and coherence.

### Dravet-prescribed ASMs modulate background ECoG activity and spectral properties in DS mice

Thermally-induced seizures are a hallmark phenotype of Dravet, and multiple studies examined the effect of ASMs on the temperature threshold of these seizures ^19,23^. However, this measurement provides a unidimensional description of a transition into a convulsive seizure and cannot be adopted for clinical use. We reasoned that analysis of the properties of background oscillations might provide additional insights into the effect of ASMs in Dravet. From these recordings, we also examined the effect of these drugs on epilepsy, quantifying the change in the frequency of interictal epileptic spikes. Interictal spikes reflect synchronous neuronal firing. They are strongly associated with epilepsy and were shown to have adverse effects on brain development and contribute to cognitive impairment ^24^. Although the relationship between Interictal spikes and spontaneous seizures is not clear ^25^, quantification of their frequency is commonly used to evaluate the severity of epilepsy and drug response in animal models. We focused on the acute effect of ASMs due to challenges with long-term ECoG recording and prolonged drug administration in immature mice.

Valproic acid (VA) is a broad-spectrum antiseizure medication and the drug of choice as a first-line treatment for Dravet. Specifically, VA was shown to reduce the frequency of spontaneous seizures in 50-70% of patients ^19,26^. In DS mice, chronic treatment with VA protected from spontaneous convulsive seizures ^19^, but its acute effects on background ECoG activity were not assessed.

When given acutely, VA has several molecular targets: *i*) increasing GABA level by reducing its degradation via inhibition of GABA transaminase and succinic semi-aldehyde dehydrogenase; *ii*) potentiation of the activity of GABA_A_ receptors; *iii*) direct inhibition of voltage-gated sodium channels and T-type voltage-gated calcium channels; *iv*) inhibition of NMDAR ^27^. In addition, long-term VA administration, which was not tested here, results in histone deacetylase inhibition, positive modulation of M currents, and activation of the ERK and JNK pathways.

First, we tested the effect of VA on the threshold of heat-induced seizures. As shown in Figure 2A, mice treated with VA (300 mg/kg) tended to convulse at higher temperatures, but this difference did not reach statistical significance. Previous results reported similar observations using the same mouse model (*Scn1a*^A1783V^ on the pure C57BL/6J background) ^23^. Conversely, others that used DS mice with a mixed genetic background demonstrated a significantly increased seizure threshold ^19,28^. Therefore, we repeated these experiments in *Scn1a*^A1783V^ mice on a mixed C57BL/6J:129x1/SvJ background. In these mice, VA significantly reduced the susceptibility to thermal seizures, increasing the average threshold temperature by 1.69 ± 0.72 °C (Figure S1). Thus, genetic background affects the ability of VA to protect from febrile seizures. However, as DS mice on a mixed background were reported before to have milder epileptic phenotypes ^6,29,30^, the severely affected DS mice on the pure C57BL/6J background were used for the rest of the study.

**Figure. 2.**
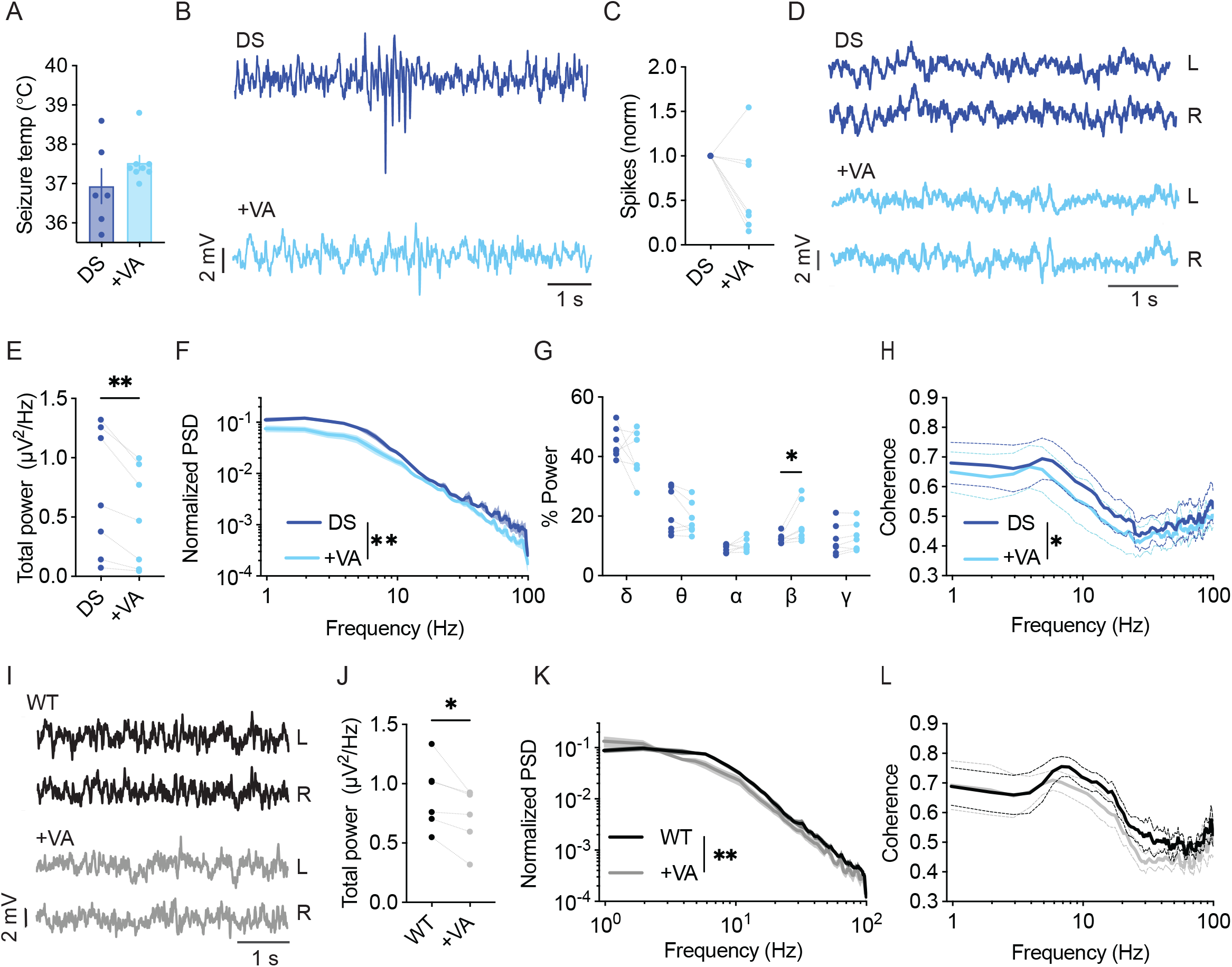
VA modulates the power and spectral properties of background ECoG. **(A)** Thermally induced seizures in mice treated with vehicle or VA (300 mg/kg). DS mice injected with saline as the vehicle, n =6; DS mice injected with VA, n = 8. **(B**) Representative traces of ECoGs from DS mice depicting epileptic activity before and after VA administration. (**C**) The change in interictal spike frequency. **(D**) Representative traces of background activity. (**E**) Absolute total power. **(F)** ECoG power density profiles normalized to the total power prior to VA administration. (**G**) Relative power in each frequency band, before and after VA. (**H**) Interhemispheric coherence plotted over the 1-100 Hz spectrum. DS, n = 7. **(I**) Representative traces of background ECoGs from WT mice before and after VA administration. **(J**) The effect of VA on total power. **(K**) ECoG power density profiles normalized to the total power prior to drug administration. (**L)** Interhemispheric coherence plotted over the 1-100 Hz spectrum. WT, n = 5. *p < 0.05, **p < 0.01.

In a different cohort of mice on the C57BL/6J background, we tested the acute effect of VA on the frequency of interictal spikes and spectral properties of background ECoG. VA had a variable impact on spike frequency, reducing the frequency by over 50% in 57% of the tested mice, with a smaller effect in 28% of the mice, and increased spike frequency in one mouse (Figure 2B-C). Focusing on quantitative analyses of background activity, VA reduced the total power spectral density (PSD) in all the tested DS mice (Figure 2D-F), in agreement with previous reports in humans ^14,15^. When inspecting the relative power of each frequency band, we observed an increase in the contribution of the beta band (Figure 2G). Despite the global effect on ECoG power, VA did not affect the coherence patterns in DS mice in any specific band. Yet, a slight overall reduction in phase coherence was observed (Figure 2H). The acute effect of VA was not specific to DS mice, and the reduction in total power was also seen in WT mice (Figure 2I-L, Figure S2A). Together, although VA had variable effects on the frequency of interictal spikes in DS mice, it reduced the power and modulated the spectral properties of background ECoG activity.

Clobazam (CLB), frequently in combination with stiripentol (STP), is another commonly used standard treatment in Dravet ^5,31^. As a monotherapy, clobazam was reported to be effective in 25-50% of patients ^26^, while the addition of STP increased responsiveness. CLB is a long-acting 1,5-benzodiazepine that acts as a positive allosteric modulator of ionotropic GABA_A_ receptors ^32^. CLB as monotherapy in mice was shown to protect from thermally induced seizures ^19,23^. STP is often administered as an adjunct drug. It enhances the effective concentration of CLB by inhibiting CYP450 isoenzymes. In addition, STP also directly modulates inhibitory GABAergic transmission via allosteric modulation of the GABA_A_ receptors, which enhances the positive effect of CLB on these channels. Moreover, STP also inhibits voltage-gated T-type calcium channels and lactate dehydrogenase (LDH), which was shown to suppress seizures ^33^.

Notably, acute administration of CLB+STP significantly reduced the susceptibility to thermally induced seizures, elevating the average temperature of convulsive seizures by ∼ 1.4 °C (Figure 3A). The effect of this drug combination was further tested using ECoG recordings. The frequency of interictal spikes decreased by over 50% in 33% of the mice, and by 20-50% in another 33%, while no change or an increase in spike frequency was observed in the rest of the mice (Figure 3B, C). Moreover, we did not observe a significant effect on the total power of background activity in DS mice (Figure 3D-F). Nevertheless, acute administration of CLB+STP altered the relative power of the delta, theta, and beta frequency bands, reducing the delta contribution and increasing that of theta and beta (Figure 3G). The interhemispheric coherence did not change after CLB+STP acute administration (Figure 3H).

**Figure 3.**
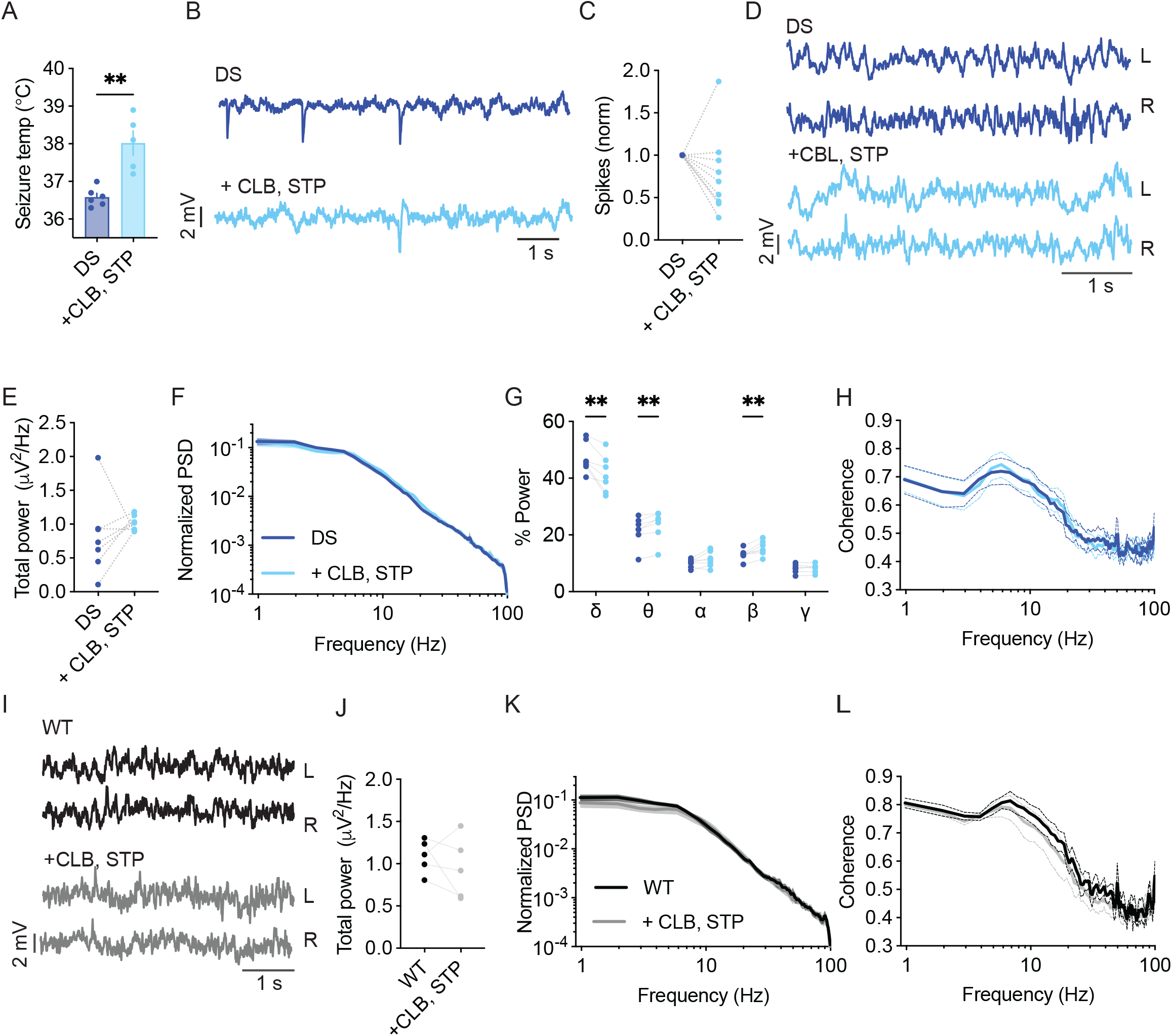
CBL+STP treatment modulates the spectral properties of background ECoG in DS mice. (**A)** CLB+STP (5,100 mg/kg, respectively) protected against hyperthermia-induced seizures in DS mice. DS mice injected with sesame oil as vehicle, n = 6; DS mice injected with CLB+STP, n = 5. (**B**) Representative traces of epileptic activity before and after acute administration of CLB+STP. (**C**) The change in the frequency of interictal spikes. (**D**) Representative examples of background activity. (**E**) The effect on total power. (**F**) ECoG power density profiles normalized to the total power prior to drug administration. (**G**) The relative power in each frequency band. (**H**) Interhemispheric coherence plotted over the 1- 100 Hz spectrum. DS, n = 7. (**I**) Representative traces of background ECoG from WT mice before and after drug administration. (**J**) The effect of CLB+STP on total power. (**K**) ECoG power density profiles normalized to the power prior to drug administration. (**L**) Interhemispheric coherence plotted over the 1-100 Hz spectrum. WT, n = 5. ** p < 0.01.

The effect of CLB+STP on WT mice was similar to that of DS mice, with no effect on the total power or coherence (Figure 3I-L), but with enhancement of the relative power of the beta frequency band (Figure S2B). Together, CLB+STP reduced the susceptibility to thermally induced seizures, reduced the frequency of interictal spikes in most of the tested mice, and modulated the spectral properties of background ECoG activity.

STP as monotherapy was shown to reduce seizure burden in patients ^19^, but it is not recommended as a monotherapy ^5^, nor was it shown to be protective against thermally induced seizures in mice ^19,23^. Nevertheless, with its direct effect on GABAergic transmission ^33^, we wondered if it would also affect the spectral properties of background ECoG activity.

STP (150 mg/kg) had variable effects on the frequency of interictal spikes, reducing their frequency by 25-53% in three mice and increasing their frequency by 35-90% in two other DS mice (Figure S3A, B). Moreover, while no significant effects were observed on the total power or coherence (Figure S3C-E, G), STP increased the contribution of the beta frequency band (Figure S3F). Together, Dravet-prescribed drugs, VA, CLB+STP, or STP alone, demonstrated variable, subject-dependent changes in the frequency of interictal epileptic spikes but with significant modulation of the spectral properties of background activity, and a common increase in the contribution of the beta frequency band.

### Contraindicated sodium channel blockers do not modulate the spectral parameters of background activity in DS mice

ASMs that act mainly via inhibition of voltage-gated sodium channels are contraindicated in Dravet and were shown to aggravate the epileptic phenotypes and adversely affect the cognitive outcome ^5,34^. Carbamazepine (CBZ) mainly targets voltage-gated sodium channels. It has an increased affinity to the inactivated state, resulting in a voltage or frequency-dependent inhibition of these channels. However, CBZ was also demonstrated to inhibit the N-type and L-type voltage-gated calcium channels and increase the secretion of serotonin ^35^.

CBZ was reported to exacerbate the susceptibility to thermally induced seizures in mice ^23^, but its effect on ECoG and background activity was not reported. Here, among the mice treated with CBZ (20 mg/kg), three out of the four mice demonstrated an increased frequency of epileptic spikes by 2-9 folds, and a marginal (3%) decrease in spike frequency was seen in the fourth mouse (Figure 4A, B). Background spectral properties analysis showed that CBZ did not modify the power, spectral profiles, or coherence (Figure 4C-G). Similarly, in WT mice, CBZ did not modify the properties of background oscillations (Figure 4H-K).

**Figure. 4.**
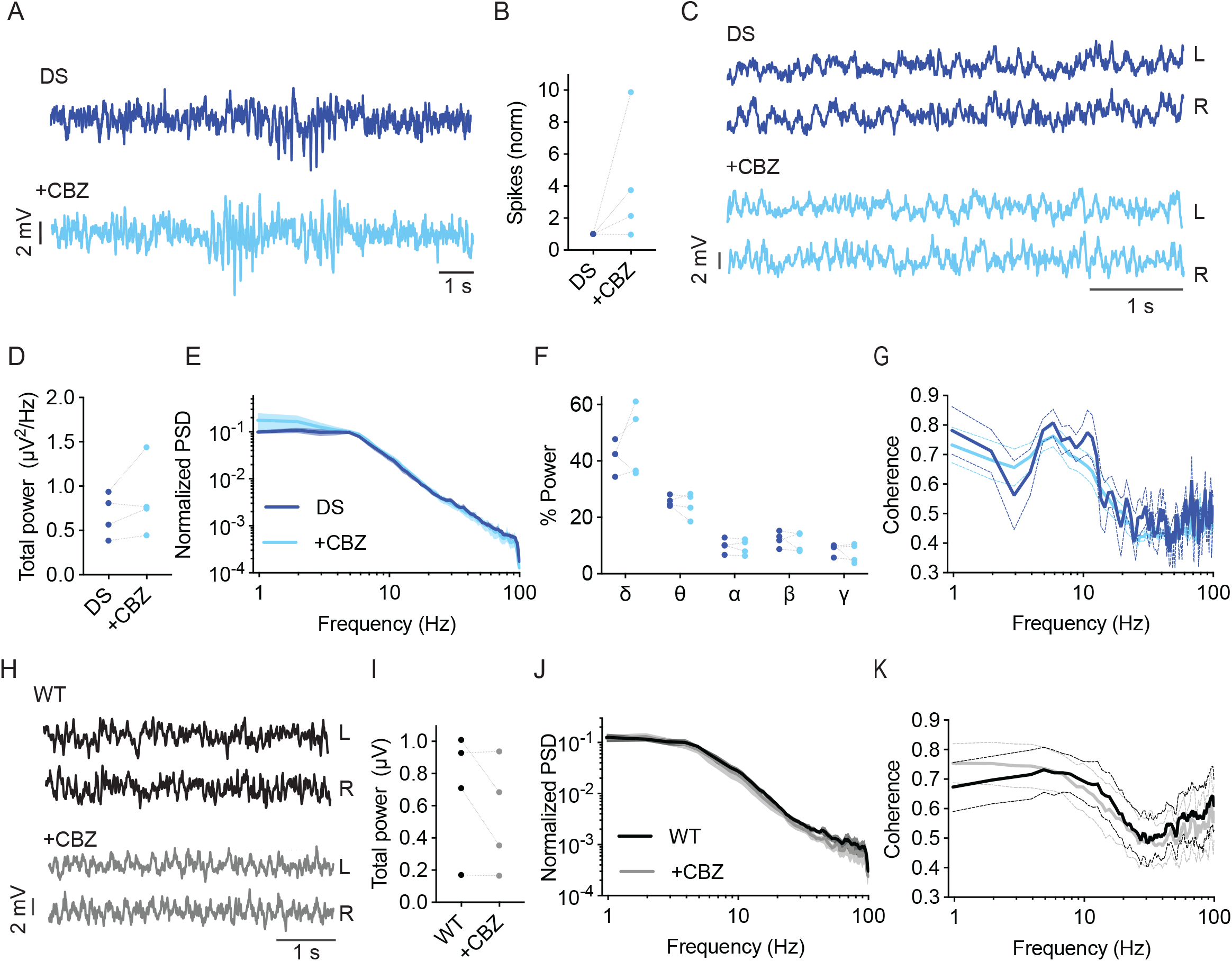
CBZ increased the frequency of interictal spikes in most DS mice. (**A)** Representative traces of epileptic activity in DS mice before and after acute administration of CBZ (20 mg/kg). (**B**) The effect of CBZ on interictal spike frequency. (**C**) Representative traces of background ECoG in DS mice before and after treatment with CBZ. **(D**) The effect on total power. (**E**) ECoG power density profiles normalized to the absolute total power prior to drug administration. (**F**) The relative power in each frequency band. (**G**) Interhemispheric coherence plotted over the 1-100 Hz spectrum. DS, n = 4. (**H**) Representative traces of background ECoG from WT mice, before and after administration of CBZ. (**I**) The effect of CBZ on total power. (**J**) ECoG power density profiles normalized to the absolute power prior to drug administration. (**K**) Interhemispheric coherence plotted over the 1-100 Hz spectrum. WT, n = 4.

Lamotrigine (LTG) is another voltage-gated sodium channel blocker avoided in Dravet ^5^. Additional mechanism of action include inhibition of voltage-gated calcium channels and reduced presynaptic glutamate release ^36^. In DS mice, acute administration of LTG (10 mg/kg) resulted in increased interictal spike frequency (Figure 5A, B). Moreover, LTG did not affect the power or spectral properties of background activity (Figure 5C-G). Together, acute administration of either CBZ or LTG increased the frequency of interictal spikes in DS mice without affecting the properties of background oscillations.

**Figure. 5.**
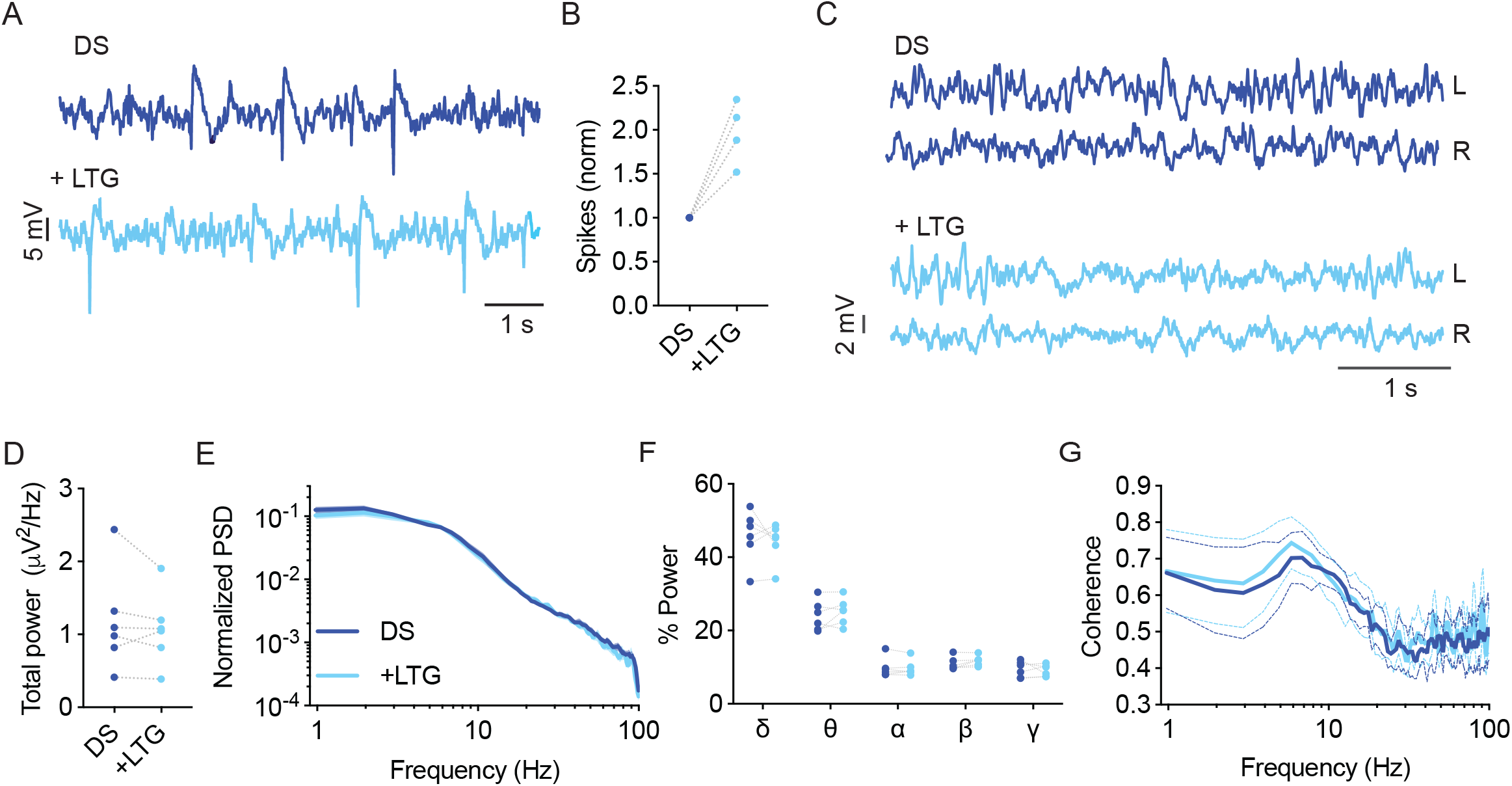
LTG increased the frequency of interictal spikes in DS mice. (**A**) Representative traces of epileptic activity in DS mice before and after acute administration of LTG (10 mg/kg). (**B**) The effect of LTG on interictal spike frequency. (**C**) Representative traces of background ECoG in DS mice before and after treatment with LTG. (**D**) The effect on total power. (**E**) ECoG power density profiles normalized to the absolute total power prior to drug administration. **(F**) The relative power in each frequency band. (G) Interhemispheric coherence plotted over the 1-100 Hz spectrum. DS, n =6.

### Correlation between interictal spike frequency, ECoG power, and spectral properties

Next, we wondered if a comprehensive analysis of the various drug-induced changes would highlight parameters that can differentiate between Dravet-prescribed and contraindicated drugs and correlate with the effect on the epileptic phenotypes as measured here by the frequency of interictal spikes.

First, we quantified the overall effect of ASMs on spike frequency. VA or CBL+STP reduced the spike frequency in 87.5% of the mice (Figures 2, 3), while STP lowered their frequency by 66% (Figure S3). Overall, a reduction of over 50% in spike frequency was achieved in ∼40% of the mice treated with Dravet-recommended ASMs. Conversely, contraindicated drugs increased the spike frequency by more than 50% in 87.5% of the tested mice (Figure 6A, B).

**Figure. 6.**
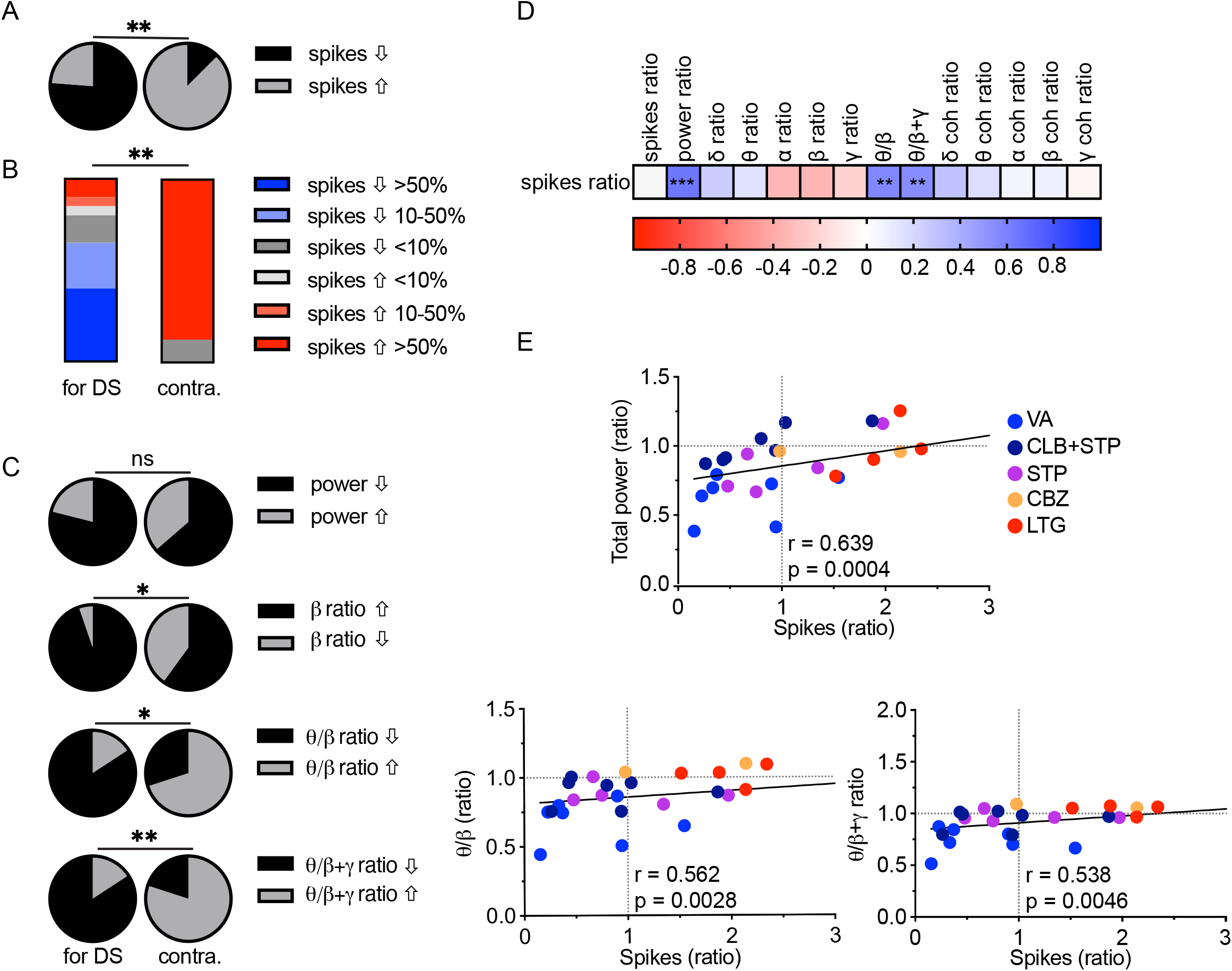
Correlations between spike frequency and spectral parameters in DS mice. **(A**) The proportion of mice with increased or decreased spike frequency among prescribed (left, VA, CLB+STP, STP) and contraindicated (right, CBZ, LTG) ASMs in DS mice. (**B**) Percentages of mice and their change in spike frequency following treatment with prescribed (left) and contraindicated (right) ASMs. (**C**) The proportion of DS mice, treated with prescribed (left) and contraindicated (right) ASMs with increased or decreased total power (top), beta contribution, theta /beta relative power ratios, or theta /(beta+gamma) ratios (bottom). (**D**) Correlation matrix (Spearman correlation) between the change in the frequency of interictal spikes, background spectral parameters, and the coherence (coh). The change was calculated as the ratio between a specific parameter after ASM administration divided same parameter before ASMs. Full correlation analysis is presented in Figure S5. A full description of the correlation coefficients and statistical significance are presented in Table S1. (**E**) Correlations between the changes in spike frequency and the total power and the (top), the theta/beta ratio (bottom, left), and the theta/(beta+gamma) ratio (bottom, right). The text indicates the Spearman correlation coefficient and statistical significance. The solid black line depicts the fit. * p < 0.05, ** p < 0.01,*** p < 0.001.

Next, we examined the relationship between background spectral changes and the effect on interictal spikes (see Figure S5 for the complete correlation analysis of spectral changes). Focusing on the total power shift, we did not see a statistical difference between the effect of Dravet-recommended and contraindicated drugs (Figure 6C). However, interestingly, the level of total power reduction correlated with a decrease in spike frequency (Figure 6D, E). This correlation was also significant for the cohort of mice treated with CLB+STP (Figure S4). Examination of the spectral power redistribution showed an increase in the relative beta contribution among the majority of mice treated with Dravet-prescribed ASMs and a less profound change after CBZ and LTG administration (Figure 6C). Nevertheless, the relative drug-induced change in the beta band contribution did not correlate with the reduction in spike frequency (Figure 6D). Thus, we wondered if power redistribution across multiple bands would correlate with the effect on interictal spikes. The ratio between theta and beta has been extensively investigated in relation to attention processes and is FDA approved for clinical confirmation of ADHD ^37^. Interestingly, a reduction in this ratio correlated with the reduction in spike frequency. A similar trend was observed when we calculated the ratio of theta to beta and gamma combined (Figure. 6 C-E). Together, examination of multidimensional readouts for the effect of acute antiseizure treatment on background activity and epileptic interictal spikes revealed a positive correlation between the frequency of interictal spikes, background power, and the relative contribution of theta beta and gamma frequency bands.

## Discussion

Dravet syndrome is characterized by multiple neuronal and circuitry changes ^10^. Here, by examining background activity in DS mice and their WT littermates, we show attenuated power of background activity in DS mice, similar to previous reports ^12,13^, and add a reduction in interhemispheric phase coherence (Figure 1). Although epileptic activity is characterized by hypersynchronous firing ^38^, seizure onset was shown to be associated with reduced synchronization ^39^, and transition to thermally induced seizures in DS mice was associated with decreased synchronization of excitatory and inhibitory neurons ^40^. In Alzheimer’s (AD) patients, reduced neural synchrony was associated with AD-related cognitive deficits ^41^. Therefore, it is possible that decreased background synchronization is also related to developmental delay and cognitive deficits in Dravet.

Interestingly, none of the tested ASMs corrected the attenuated power or coherence (Figures 1-3, S3)., and a reduction in total power correlated with a decrease in the frequency of interictal spikes (Figure 6). However, as low power was suggested as a risk factor for premature mortality in DS mice ^12,13^, an ideal treatment for Dravet can be expected to amend these parameters rather than further decrease the power. Thus, it may be that the observed association between reduced power and reduction in spike frequency provides an electrographic measurement for the partial ability of ASMs to correct Dravet, stressing the need for developing novel treatment options for Dravet.

Examination of the power redistribution following acute drug administration, demonstrated that Dravet-prescribed ASMs modulate the spectral properties of background activity, mainly by affecting the contribution of the beta frequency band. In accordance, increased beta power was observed in patients on benzodiazepines or barbiturates^42^, and acute administration of VA ^14^ or CLB ^16^ was shown to increase the contribution of beta. Moreover, increased beta power was observed in medicated Dravet patients at the age of 1-2 years compared to unmedicated age match controls ^3^. Of note, this age represents the onset of spontaneous seizures, which is somewhat parallel the development stage of the mice used here. Furthermore, an increase in beta contribution was also suggested to predict the positive effect of cannabidiol as an add-on therapy in patients treated with CLB ^42^. Despite that, in our cohort, the relative change in beta band contribution alone did not correlate with the reduction in spike frequency (Figure 6C, D). However, a broader examination of the power redistribution between theta, beta, and gamma bands, demonstrated a correlation with a reduction in spike frequency (Figure 6C-E). Interestingly, the ratio between theta and beta was suggested to be a reliable electrographic measurement that reflects cognitive processing and was therefore indicated as a biomarker in patients with attention deficit disorder ^37^. Thus, we propose that examining these background activity changes following the administration of ASMs may be an additional tool to evaluate the therapeutic potential of drug treatment also in Dravet.

One limitation of the current study is that the effect on epilepsy was quantified as the change in the frequency of the interictal spikes using short-term ECoG recordings, due to challenges with prolonged recordings in immature mice. Conversely, in clinical settings, drug treatment is focused on seizure reduction and prevention of status epilepticus over a long time. Nevertheless, despite the apparent difference in readout (interictal spikes vs. spontaneous seizures) and drug regimen (acute vs. chronic), the overall response levels reported here resembled clinical reports ^19,26,43^. Specifically, patients show 50-80% responsiveness levels, compared to a reduction in spike frequency in ∼80% of the tested mice (Figures 2, 3, 6). In contrast, drugs known to aggravate Dravet epileptic phenotypes increased the frequency of spikes (Figures 4-6). Interestingly, even though we used DS mice of the same age, *Scn1a* mutation (A1783V) and C57BL/6J genetic background, we still observed variable responses. This may indicate that distinct individual epileptogenic processes and various homeostatic changes may contribute to personalized drug responses.

Together, this study focused on measurements of background properties and interictal spikes in mice. Some of these parameters, like the lower interhemispheric coherence compared to WT mice, or measurements of the absolute power that are sensitive to the equipment used, recording quality, and the developmental stage, are more suitable for pre-clinical studies than for clinical use. Conversely, we propose that alteration in the spectral properties, specifically the correlation between the ratio of theta to beta and gamma, can potentially be applied in patients (Figure 6). Nevertheless, as our analyses are based on the treatment of naïve mice, it remains to be seen if a stepwise approach in which ASMs are given acutely in conjunction with clinical EEG monitoring will result in similar spectral changes in patients.

In conclusion, we describe the acute effects of ASMs on ECoG spectral properties in DS mice. Our analyses, demonstrate a distinct impact of Dravet-prescribed drugs on ECoG spectral properties and interictal spikes frequency, suggesting that such measurements can potentially be used as additional informative readouts for the therapeutic benefit of drug therapy in Dravet.

## Acknowledgment

This study was supported by the Israel Science Foundation (grant 1454/17, MR). Support also came from The Claire and Amedee Maratier Institute for the Study of Blindness and Visual Disorders, Sackler Faculty of Medicine, Tel-Aviv University (MR), and The Stolz Foundation Sackler Faculty of Medicine, Tel-Aviv University (MR). Support also came from ERA-NET E-Rare (MR) and The American Dravet Syndrome Foundation (MR).

## Author Contributions

This work was performed in partial fulfillment of the requirements for a Ph.D. degree of SQ. SQ, MB, and MR conceived the project, designed the experiments, and analyzed the data. MB and SQ performed the experiments, SQ and MR and wrote the manuscript. SQ and MO developed the analysis codes. All authors contributed to the article and approved the submitted version.

