## Supplementary Materials: Supplementary methods, Figures S1-S5, Table S1 for "Acute effect of antiseizure drugs on background oscillations in *Scn1a*^A1783V^ Dravet syndrome mouse model"

#### Animals and Surgery

DS mice harboring the global *Scn1a*$^{A1783V/WT}$ mutation on pure C57BL/6J background were generated as described before [1,2], by crossing conditional *Scn1a*$^{A1783V}$ males (The Jackson Laboratory, stock #026133) with CMV-Cre females (The Jackson Laboratory, stock #006054). Electrode implantation was done on P19-P25, as previously described [1]. Briefly, the mice were anesthetized (ketamine/xylazine, 191/4.25 mg/kg in normal saline) by IP injection. For analgesia, carprofen (5 mg/kg) was injected prior to surgery and 24 hours post-op. A midline incision was made above the skull, and fine silver wire electrodes (130 µm diameter bare; 180 µm diameter coated) were placed at visually identified locations, bilaterally above the somatosensory cortex; a reference electrode was placed on the cerebellum midline; ground and electromyography (EMG) electrodes were placed subcutaneously over the left and right shoulders, respectively. The electrodes were secured using dental cement and connected to a micro-connector system. Mice were allowed to recover for at least 48 hours before recording.

#### Data acquisition and ECoG signal pre-processing

Video-ECoG recordings were obtained from freely behaving mice at P21-P27, as described before [1]. The recordings lasted for 4 hours, between 10 am and 5 pm. After two hours of recording, one of the drugs was administered. The first 30 minutes of the recording and the first 30 minutes post-drug administration were considered acclimation periods and were not taken for analysis. The EMG signal was denoised and smoothed using a custom-written Python script, based on methods described before [3]. Briefly, the raw EMG signal was denoised with the Teager-Kaiser Energy Operator (TKEO), and the output was rectified and smoothed across a 3Hz cut-off low pass filter. Next, to calculate spectral properties, the recording was divided into 5 seconds epochs with a 2.5-second sliding window. Power spectral densities of the ECoG were computed for each epoch using Welch's method with a 50% overlap Hann window. We used a custom-written threshold algorithm in python to extract artifact-free epochs where the mice were awake and not moving. An epoch was taken for analysis if all conditions were met: *i*) the delta power (1-4Hz) of the epoch was lower than the median delta power; *ii*) the theta (4-8Hz) to delta ratio was lower than the 75% quantile of the entire 1.5 hours recording block; *iii*) the enveloped EMG was less than two times the harmonic mean throughout the entire epoch [4,5]. Finally, all automatically selected epochs were concatenated and manually inspected. For each mouse, at least 100 epochs were analyzed. Then, the data were averaged across the different epochs. Thus, only one data point before drug administration, and one data point after ASM administration, was considered for statistical analysis. To quantify phase synchrony between the left and right somatosensory electrodes we calculated the interhemispheric coherence coefficient (coherence) using a custom-written Python script, based on the coherence coefficient equation as described before [6].

Custom-written codes are available at https://github.com/shirquinn/Rubinstein_Lab.git.

To quantify the frequency of interictal spikes, we applied a bandpass filter between 0.5 and 60 Hz and used the spike histogram module in LabChart 8 software (ADInstruments). The threshold was set to 4-5 times the standard deviation of the filtered signal, with a 50 ms pretrigger interval and 250 ms maximal total duration. The spikes were then manually inspected in the scope view window, and artifacts were rejected.

### Statistics

Statistical analyses were performed using GraphPad Prism 9 software (GraphPad Software). Data are reported as mean ± SE, and the individual data points are depicted on the bar graphs. For comparison between wild-type (WT) and DS mice and the effect of drugs on thermal induced seizures, we used the Student's t-test (for data with normal distribution) or the non-parametric Mann-Whitney test (for data that did not distribute normally). To compare WT or DS mice before and after drug administration, we used the parametric paired Student's t-test (for data with normal distribution), or the Wilcoxon matched-pairs signed rank test (for data that did not distribute normally). To compare the effect on normalized power spectral density (PSD) or coherence, we used Two-Way repeated measures ANOVA followed by Sidak posthoc analysis. For the correlation analysis, the Spearman rank-order correlation coefficient was used. Statistical analysis of pie charts was performed using Fisher's exact test. We considered $p < 0.05$ as statistically significant.

### Supplementary Figures:

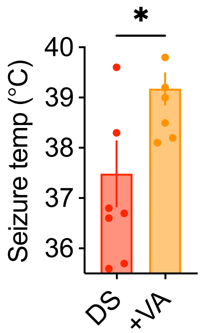


**Figure S1. Valproic acid (300 mg/kg) decreased the susceptibility of thermally induced seizures in *Scn1a*$^{A1783V/WT}$ mice on a mixed background (50:50 C57BL/6J:129x1/SvJ).**

The mice were generated by crossing *Scn1a*$^{A1783V/WT}$ males (pure C57BL/6J) with WT females (129x1/SvJ). DS injected with saline n=7; DS injected with VA n=6.

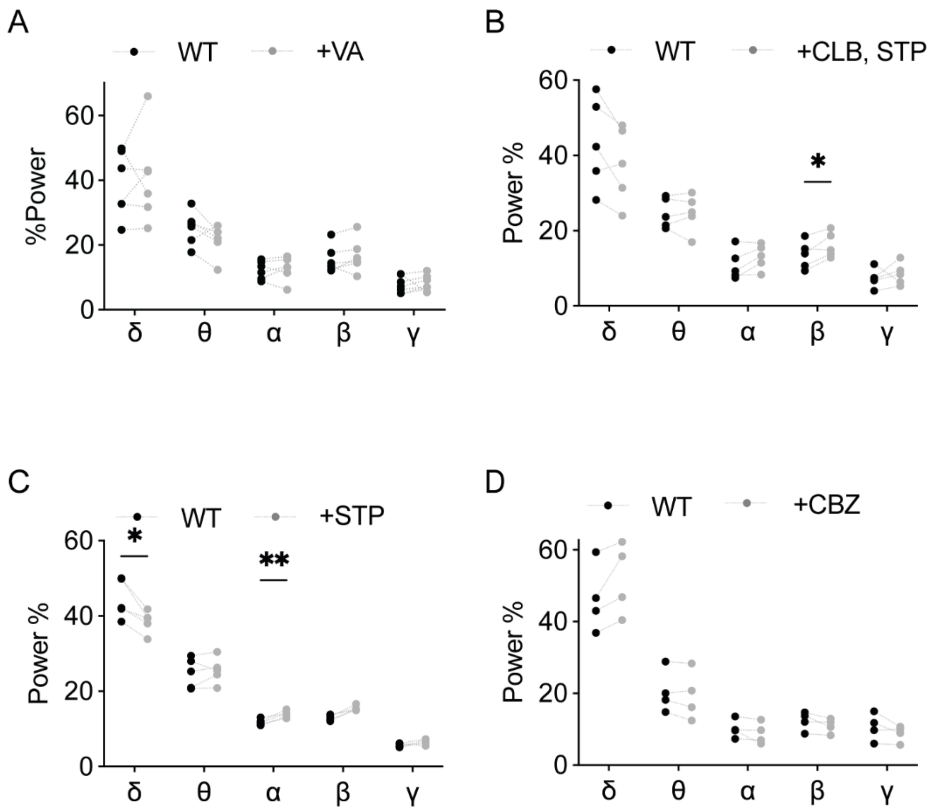


**Figure S2. The effect of ASMs on spectral properties of WT mice.**

**(A)** Relative power in each frequency band, before and after administration of VA, related to Figure 2I-K. n = 6. **(B)** Relative power in each frequency band, before and after administration of CLB+STP, related to Figure 3I-K. n = 5. **(C)** Relative power in each frequency band, before and after administration of STP, related to Figure S3H-KJ. n = 5. **(D)** Relative power in each frequency band, before and after administration of CBZ, related to Figure 4H-J. n = 4. *$p < 0.05$, **$p < 0.01$.

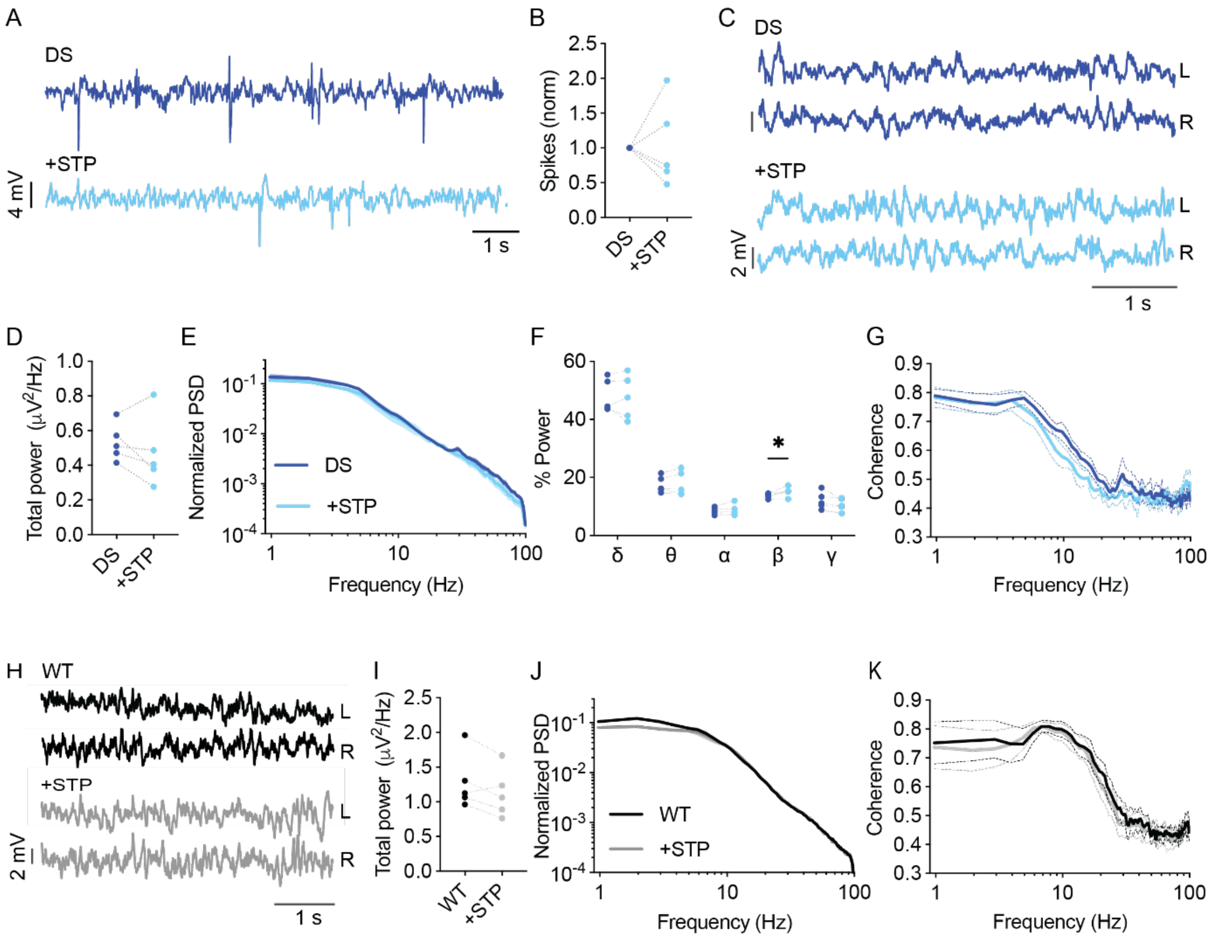


**Figure S3. Analysis of background EcoG activity in response to STP administration.**

**(A)** Representative traces of epileptic activity in DS mice prior to and post, acute administration of STP (150 mg/kg). **(B)** The effect of STP on interictal spike frequency. **(C)** Representative traces of background ECoG in DS mice before and after STP administration. **(D)** The effect on total power. **(E)** ECoG power density profiles normalized to the absolute total power prior to drug administration. **(F)** The relative power in each frequency band. **(G)** Interhemispheric coherence plotted over the 1- 100 Hz spectrum. DS, n = 5. **(H)** Representative traces of background ECoG. From WT mice, before and after drug administration. **(I)** The effect of STP on total power. **(J)** ECoG power density profiles normalized to the absolute power prior to drug administration. **(K)** Interhemispheric coherence plotted over the 1- 100 Hz spectrum. WT, n = 5. * $p < 0.05$.

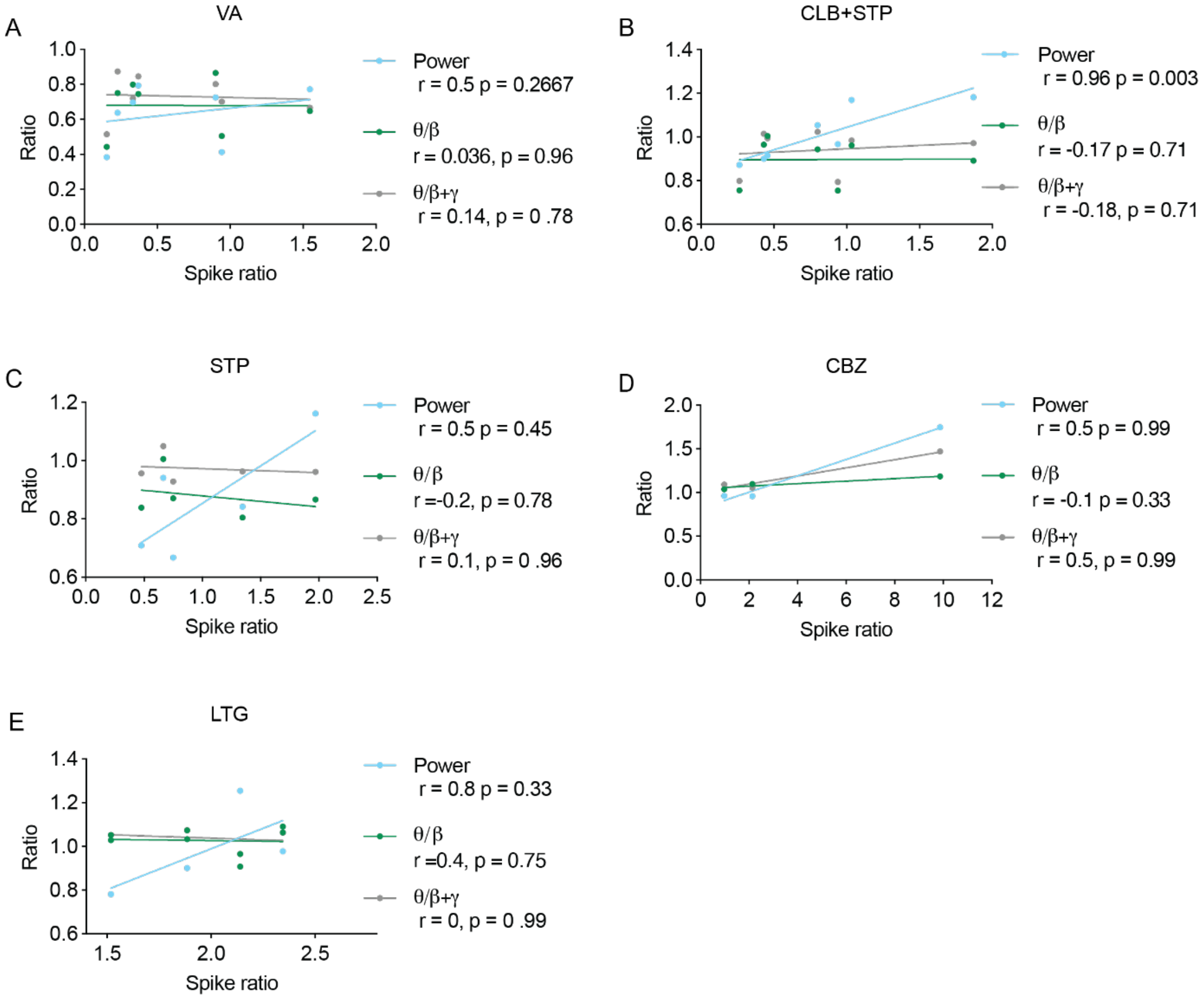


**Figure S4. Correlations between spike frequency and spectral parameters in DS mice**.

Correlation between the change in total power, theta /beta ratios, and theta /(beta + gamma) ratios in DS mice treated with **(A)** VA, n = 7. **(B)** CLB+STP, n = 9. **(C)** STP, n = 5. **(D)** CBZ, n = 3. (E) LTG, n = 4. The correlation coefficients (Spearman, r) and p-values (p) are depicted. The solid line is the result of simple linear regression and is depicted for illustration purposes. Related to Figure 6E.

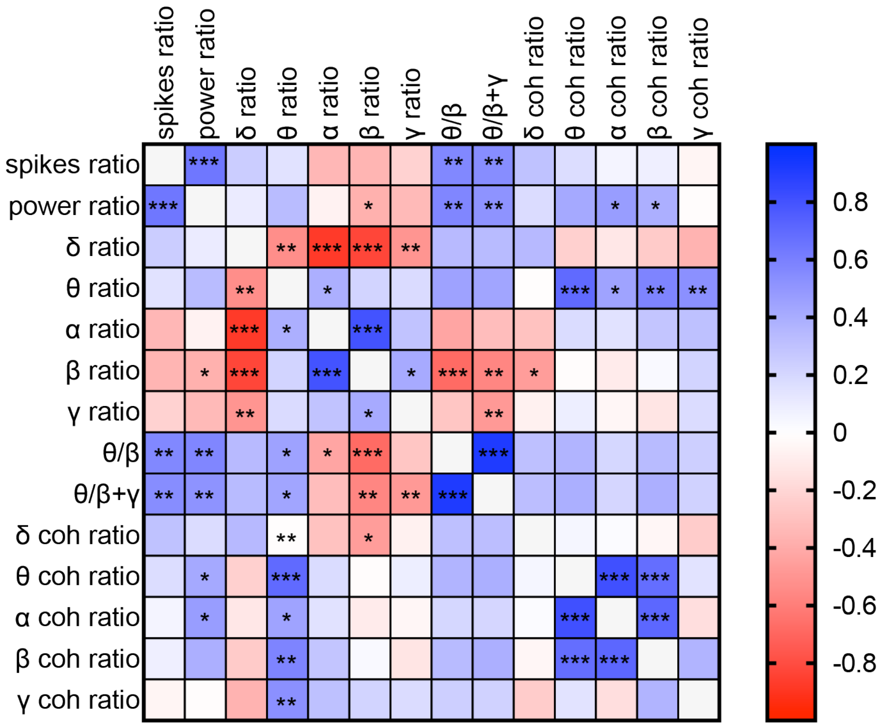


**Figure S5. Correlations between spike frequency and spectral parameters in DS mice**.

The full Correlation matrix (Spearman correlation) between the change in the frequency of interictal spikes, background spectral parameters and the coherence (coh). A full description of the correlation coefficients and statistical significance are presented in Table S1.

### Table S1 – **Spearman correlation coefficient and p values**

Spearman r

| | spikes ratio | power ratio | δ ratio | θ ratio | α ratio | β ratio | γ ratio | θ/β | θ/β+γ | δ coh ratio | θ coh ratio | α coh ratio | β coh ratio | γ coh ratio |
|---|---|---|---|---|---|---|---|---|---|---|---|---|---|---|
| **spikes ratio** | 1.000 | 0.640 | 0.240 | 0.138 | -0.342 | -0.354 | -0.216 | 0.562 | 0.538 | 0.296 | 0.162 | 0.048 | 0.073 | -0.051 |
| **power ratio** | 0.640 | 1.000 | 0.094 | 0.324 | -0.061 | -0.375 | -0.330 | 0.571 | 0.510 | 0.166 | 0.412 | 0.469 | 0.382 | -0.008 |
| **δ ratio** | 0.240 | 0.094 | 1.000 | -0.528 | -0.876 | -0.829 | -0.500 | 0.331 | 0.326 | 0.334 | -0.228 | -0.116 | -0.255 | -0.363 |
| **θ ratio** | 0.138 | 0.324 | -0.528 | 1.000 | 0.381 | 0.208 | 0.175 | 0.446 | 0.430 | -0.007 | 0.694 | 0.440 | 0.577 | 0.525 |
| **α ratio** | -0.342 | -0.061 | -0.876 | 0.381 | 1.000 | 0.798 | 0.288 | -0.424 | -0.317 | -0.294 | 0.172 | 0.138 | 0.279 | 0.298 |
| **β ratio** | -0.354 | -0.375 | -0.829 | 0.208 | 0.798 | 1.000 | 0.406 | -0.681 | -0.563 | -0.464 | -0.010 | -0.095 | 0.021 | 0.210 |
| **γ ratio** | -0.216 | -0.330 | -0.500 | 0.175 | 0.288 | 0.406 | 1.000 | -0.275 | -0.482 | -0.067 | 0.079 | -0.041 | -0.127 | 0.166 |
| **θ/β** | 0.562 | 0.571 | 0.331 | 0.446 | -0.424 | -0.681 | -0.275 | 1.000 | 0.910 | 0.305 | 0.362 | 0.196 | 0.326 | 0.233 |
| **θ/β+γ** | 0.538 | 0.510 | 0.326 | 0.430 | -0.317 | -0.563 | -0.482 | 0.910 | 1.000 | 0.311 | 0.386 | 0.198 | 0.382 | 0.216 |
| **δ coh ratio** | 0.296 | 0.166 | 0.334 | -0.007 | -0.294 | -0.464 | -0.067 | 0.305 | 0.311 | 1.000 | 0.043 | 0.013 | -0.042 | -0.245 |
| **θ coh ratio** | 0.162 | 0.412 | -0.228 | 0.694 | 0.172 | -0.010 | 0.079 | 0.362 | 0.386 | 0.043 | 1.000 | 0.810 | 0.681 | 0.137 |
| **α coh ratio** | 0.048 | 0.469 | -0.116 | 0.440 | 0.138 | -0.095 | -0.041 | 0.196 | 0.198 | 0.013 | 0.810 | 1.000 | 0.713 | -0.158 |
| **β coh ratio** | 0.073 | 0.382 | -0.255 | 0.577 | 0.279 | 0.021 | -0.127 | 0.326 | 0.382 | -0.042 | 0.681 | 0.713 | 1.000 | 0.358 |
| **γ coh ratio** | -0.051 | -0.008 | -0.363 | 0.525 | 0.298 | 0.210 | 0.166 | 0.233 | 0.216 | -0.245 | 0.137 | -0.158 | 0.358 | 1.000 |

### P values

| | spikes ratio | power ratio | δ ratio | θ ratio | α ratio | β ratio | γ ratio | θ/β | θ/β+γ | δ coh ratio | θ coh ratio | α coh ratio | β coh ratio | γ coh ratio |
|---|---|---|---|---|---|---|---|---|---|---|---|---|---|---|
| **spikes ratio** | | 0.0004 | 0.2383 | 0.5000 | 0.0870 | 0.0762 | 0.2883 | 0.0028 | 0.0046 | 0.1506 | 0.4382 | 0.8181 | 0.7285 | 0.8096 |
| **power ratio** | 0.0004 | | 0.6273 | 0.0868 | 0.7529 | 0.0451 | 0.0804 | 0.0012 | 0.0047 | 0.4163 | 0.0365 | 0.0157 | 0.0542 | 0.9696 |
| **δ ratio** | 0.2383 | 0.6273 | | 0.0033 | 0.0000 | 0.0000 | 0.0057 | 0.0799 | 0.0848 | 0.0954 | 0.2625 | 0.5729 | 0.2092 | 0.0680 |
| **θ ratio** | 0.5000 | 0.0868 | 0.0033 | | 0.0413 | 0.2780 | 0.3642 | 0.0152 | 0.0200 | 0.9722 | 0.0001 | 0.0245 | 0.0020 | 0.0059 |
| **α ratio** | 0.0870 | 0.7529 | 0.0000 | 0.0413 | | 0.0000 | 0.1295 | 0.0220 | 0.0936 | 0.1444 | 0.4009 | 0.5021 | 0.1681 | 0.1386 |
| **β ratio** | 0.0762 | 0.0451 | 0.0000 | 0.2780 | 0.0000 | | 0.0289 | 0.0000 | 0.0015 | 0.0170 | 0.9617 | 0.6454 | 0.9195 | 0.3026 |
| **γ ratio** | 0.2883 | 0.0804 | 0.0057 | 0.3642 | 0.1295 | 0.0289 | | 0.1482 | 0.0081 | 0.7463 | 0.7014 | 0.8410 | 0.5369 | 0.4163 |
| **θ/β** | 0.0028 | 0.0012 | 0.0799 | 0.0152 | 0.0220 | 0.0000 | 0.1482 | | 0.0000 | 0.1303 | 0.0691 | 0.3375 | 0.1035 | 0.2524 |
| **θ/β+γ** | 0.0046 | 0.0047 | 0.0848 | 0.0200 | 0.0936 | 0.0015 | 0.0081 | 0.0000 | | 0.1223 | 0.0515 | 0.3324 | 0.0542 | 0.2899 |
| **δ coh ratio** | 0.1506 | 0.4163 | 0.0954 | 0.9722 | 0.1444 | 0.0170 | 0.7463 | 0.1303 | 0.1223 | | 0.8358 | 0.9485 | 0.8384 | 0.2274 |
| **θ coh ratio** | 0.4382 | 0.0365 | 0.2625 | 0.0001 | 0.4009 | 0.9617 | 0.7014 | 0.0691 | 0.0515 | 0.8358 | | 0.0000 | 0.0001 | 0.5042 |
| **α coh ratio** | 0.8181 | 0.0157 | 0.5729 | 0.0245 | 0.5021 | 0.6454 | 0.8410 | 0.3375 | 0.3324 | 0.9485 | 0.0000 | | 0.0000 | 0.4419 |
| **β coh ratio** | 0.7285 | 0.0542 | 0.2092 | 0.0020 | 0.1681 | 0.9195 | 0.5369 | 0.1035 | 0.0542 | 0.8384 | 0.0001 | 0.0000 | | 0.0726 |
| **γ coh ratio** | 0.8096 | 0.9696 | 0.0680 | 0.0059 | 0.1386 | 0.3026 | 0.4163 | 0.2524 | 0.2899 | 0.2274 | 0.5042 | 0.4419 | 0.0726 | |
